# Vasopressin regulates social play behavior in sex-specific ways through glutamate modulation in the lateral septum

**DOI:** 10.1101/2023.03.31.535148

**Authors:** Remco Bredewold, Catherine Washington, Alexa H. Veenema

**Affiliations:** Neurobiology of Social Behavior Laboratory, Department of Psychology and Neuroscience Program, Michigan State University, East Lansing, MI, USA

## Abstract

Social play is a highly rewarding behavior that is essential for the development of social skills. Social play is impaired in children diagnosed with autism, a disorder with a strong sex bias in prevalence. We recently showed that the arginine vasopressin (AVP) system in the lateral septum (LS) regulates social play behavior sex-specifically in juvenile rats: Administration of a AVP 1a receptor (V1aR) antagonist increased social play behavior in males and decreased it in females. Here, we demonstrate that glutamate, but not GABA, is involved in the sex-specific regulation of social play by the LS-AVP system. First, males show higher extracellular glutamate concentrations in the LS than females while they show similar extracellular GABA concentrations. This resulted in a baseline sex difference in excitatory/inhibitory balance, which was eliminated by V1aR antagonist administration into the LS: V1aR antagonist increased extracellular glutamate release in females but not in males. Second, administration of the glutamate receptor agonist L-glutamic acid into the LS prevented the V1aR antagonist-induced increase in social play behavior in males while mimicking the V1aR antagonist-induced decrease in social play behavior in females. Third, administration of the glutamate receptor antagonists AP-5 and CNQX into the LS prevented the V1aR antagonist-induced decrease in social play behavior in females. Last, both sexes showed increases in extracellular LS-GABA release upon V1aR antagonist administration into the LS and decreases in social play behavior upon administration of the GABA-A receptor agonist muscimol into the LS, suggesting that GABA is not involved in the sex-specific regulation of social play by the LS-AVP system. Finally, to start identifying the cellular mechanism mediating the sex-specific effects of the LS-AVP system on LS-glutamate, we determined the presence of potential sex differences in the type of LS cells expressing V1aR. However, no sex differences were found in the percentage of Avpr1a+ LS cells expressing markers for either GABAergic neurons, somatostatin-expressing neurons, calbindin 1-expressing neurons, or astrocytes. In conclusion, these findings demonstrate that the LS-AVP system regulates social play sex-specifically via differential local glutamatergic neurotransmission in male and female juvenile rats. Further research is required to uncover the underlying cellular mechanism.

## INTRODUCTION

Social play among juvenile peers has been shown to facilitate the acquisition of social and cognitive competence in humans, non-human primates, and rats (Palagi et al., 2016; Pellegrini, 1988; Suomi & Harlow, 1972; Sigman & Ruskin, 1999; Hol et al., 1999; Guralnick et al., 2006; Cordoni & Palagi, 2011; Van Den Berg et al., 1999). Furthermore, experiments in non-human primates and rats have demonstrated that social play is a highly rewarding behavior (Achterberg et al, 2016; Calcagnetti and Schechter, 1992; Ikemoto and Panksepp, 1992; Normansell and Panksepp, 1990; Trezza et al, 2009). Children diagnosed with autism spectrum disorder (ASD) show deficits in social play (Mundy, 1995; Jordan, 2003) which are likely due to disrupted reward-seeking tendencies, especially in social contexts (Chevallier et al., 2012; Kohls et al., 2012, 2013). Moreover, ASD shows robust sex differences in incidence, symptoms severity, and treatment responses (Rutter et al., 2003). Together, this indicates that it is essential to uncover the neurobiological underpinnings of the naturally rewarding aspects of social play and its regulation in both sexes with the goal to inform therapeutic strategies aimed at restoring social play deficits.

The lateral septum (LS) is a key brain region involved in social behavior, motivation, and reward (Clarke and File, 1982; Goodson et al., 1997; Beiderbeck et al., 2007; Scotti et al., 2011; Luo et al., 2011; McDonald et al., 2012; Veenema et al., 2012; Lukas et al., 2013; Harasta et al., 2015). More specifically, the LS plays a critical role in juvenile social play behavior as demonstrated in both rats and hamsters (Beatty et al., 1982; Cheng & Delville, 2009; Veenema et al., 2013, Bredewold et al., 2014, 2015). We previously showed that arginine-vasopressin (AVP), glutamate, and GABA systems in the LS each modulate social play behavior in both male and female juvenile rats (Veenema et al., 2013, Bredewold et al., 2014, 2015). Of these three systems, AVP and glutamate regulated social play behavior in sex-specific ways. In detail, administration of the AVP V1a receptor (V1aR) antagonist into the LS increased social play levels in males and decreased it in females (Veenema et al., 2013, Bredewold et al., 2014) while administration of glutamate receptor antagonists into the LS decreased social play levels in females, while no changes in social play levels were seen in males (Bredewold et al., 2015). In vitro electrophysiological recordings of LS neurons suggest that AVP acting on the V1aR enhances excitatory responses to glutamate (Joels & Urban, 1982, 1984, 1985) and enhances GABAergic inhibitory input (Allaman-Exertier et al., 2007). The latter could be mediated directly by V1aRs expressed on GABAergic interneurons, which are the main cell type in the LS (Risold & Swanson, 1997a; Garrido Sanabria et al., 2006). Taken together, we hypothesized that AVP, acting on V1aR in the LS, regulates social play differently in males and females by sex-specific modulation of glutamate and/or GABA neurotransmission in the LS.

To study this, we first used intracerebral microdialysis and retrodialysis to measure changes in extracellular release of glutamate and GABA in the LS in response to LS-V1aR blockade and exposure to social play. Second, we used acute pharmacological manipulations to determine the causal involvement of glutamate and GABA signaling in the LS-V1aR blockade-induced sex difference in social play behavior. Last, we used fluorescent *in situ* hybridization to determine whether there are sex differences in the type of V1aR mRNA-expressing cells in the LS. This would provide a first step in identifying the cellular mechanism by which LS-V1aR activation causes the sex-specific regulation of social play behavior.

## EXPERIMENTAL PROCEDURES

### 1. Animals

Male and female Wistar rats (23 days of age) were obtained from Charles River (Raleigh, NC) and maintained under standard laboratory conditions (12 h light/dark cycle, lights off at 14:00 h, 22 °C, 50% humidity, food and water *ad libitum*). Rats were housed in same-sex groups of four in standard rat cages (48 × 27 × 20 cm) unless otherwise mentioned. The experiments were conducted in accordance with the guidelines of the NIH and approved by the Institutional Animal Care and Use Committee at Michigan State University.

### 2. Social play test

During the beginning of the dark phase (between 14:00 h and 15:00 h) and under red light conditions, each experimental rat was exposed in its home cage to an unfamiliar age- and sex-matched stimulus rat for 10 min. Experimental rats were single housed for 2-3 days while the stimulus rats were group-housed. These conditions allow testing for self-initiated social play by the experimental rat. This has translational relevance because self-initiated play is found to be impaired in ASD children (Mundy, 1995; Jordan, 2003). All tests were videotaped for subsequent analysis of behavior by a researcher blind to the treatment condition using JWatcher (http://www.jwatcher.ucla.edu/). The duration of social play was scored according to Veenema & Neumann (2009) and consisted of the total amount of time spent in playful social interactions that included nape attacks (when the experimental rat displays nose attacks or nose contacts toward the nape of the neck of the stimulus rat), pinning (when the experimental rat holds the stimulus rat on its back in a supine position), and supine poses (when the experimental rat is being pinned by the stimulus rat).

### 3. Experiment 1: Effects of LS-V1aR blockade on extracellular glutamate and GABA release in the LS under baseline conditions and during exposure to social play in male and female juvenile rats

Glutamate and GABA concentrations were measured in the LS of male and female juvenile rats before, during, and after exposure to the social play test and with or without infusion of a specific V1aR antagonist into the LS using microdialysis combined with retrodialysis.

#### 3.1. Microdialysis Probe Surgery

Juvenile male and female rats (30 days of age), were anesthetized with isoflurane (Butler Schein Animal Health, Dublin, OH) and mounted on a stereotaxic frame with the tooth bar set at −4.5 mm. The u-shaped microdialysis probes (Brainlink, Groningen, the Netherlands) were unilaterally implanted into the LS according to Lukas et al. (2011) at coordinates 0.4 mm caudal to bregma, −1.5 mm lateral to the midline, 5.6 mm beneath the surface of the skull with an angle of 10° to avoid damage to the sagittal sinus. The probes were filled with sterile Lactated Ringer’s solution (pH 7.4) and fixed to the skull with three stainless steel screws and dental cement. Two 5 cm long pieces of polyethylene tubing (PE 20, Plastics One, Roanoke, VA) filled with Ringer’s solution were connected to the inflow and the outflow of the probe and fixed with dental cement.

#### 3.2. Microdialysis and retrodialysis procedure

One day after surgery, probes were flushed with Ringer’s and rats were familiarized to the microdialysis sampling procedure. Two days after surgery, probes were connected via PE20 tubing to a syringe mounted onto a microinfusion pump and perfused with Ringer’s (3.0 µl/min, pH 7.4) to establish equilibrium between the inside and outside of the microdialysis membrane. Two hours later, six consecutive 10-min dialysates were collected per rat according to Bredewold et al. (2015) and Lukas et al (2011). In detail, dialysates 1 and 2 were taken under baseline (undisturbed) conditions. Then either Ringer’s solution (vehicle-treated, n = 8/sex) or Ringer’s solution containing 10 µg/ml of the specific V1aR antagonist d(CH_2_)_5_Tyr(Me^2^)AVP (V1aR-A; Manning et al., 2008; V1aR-A-treated males, n = 9; V1aR-A-treated females, n=10) was perfused during dialysates 3 and 4, Dialysate 4 also included rats being exposed to the 10-min social play test. Dialysates 5 and 6 were taken thereafter with V1aR-A-treated rats being switched back to Ringer’s solution at the beginning of dialysate 5. Microdialysates were collected in 0.5 ml Eppendorf tubes and were immediately frozen on dry ice and subsequently stored at -45°C until quantification for glutamate and GABA using HPLC-MS/MS. Rats were killed with CO_2_ and proper probe placement was checked histologically on Nissl-stained cryocut coronal brain sections. Rats with incorrect probe placement were removed from the analysis.

#### 3.3. Analysis of glutamate and GABA with LC-MS/MS

Concentrations of GABA and glutamate in dialysates were quantified using an integrated HPLC-MS/MS system (BrainsOnline, San Francisco, CA) according to Bredewold et al. (2015). Briefly, 10 µl dialysate samples were mixed with 10 µl 0.02 M formic acid in water containing 0.04% ascorbic and 4 µl stable labelled isotope mixture containing GABA labelled with 6 deuterium and glutamate labelled with 5 deuterium. The mixtures were then placed in an autosampler (SIL-20 HT with pre-treatment option, Shimadzu, Japan) and 30 µl of derivatization mixture, containing 5 mg/ml Symdaq reagent in 0.5M NaHCO_3_ buffer (Brainlink, the Netherlands), was automatically added by the autosampler. After 2 min reaction time, 45 µl of the mixture was injected onto the HPLC Column. Chromatographic separation was performed on a reversed phase column (100 *3.0 mm Synergi MAX-RP C12, particle size 2.5 µm, Phenomenex, Torrance, CA, USA) held on a temperature of 35° Celsius in a column oven (CTO-20, Shimadzu, Japan). Samples were eluted using a linear gradient of 0.2–70 % of acetonitrile with 0.1 % formic in ultrapure water with 0.1 % formic acid for 5.3 min at a flow rate of 0.3 ml/min (LC-20 AD, Shimadzu, Japan). The flow was mixed (postcolumn) with a makeup flow of 0.15 ml/min of the mobile phase (acetonitrile with 0.1 % formic acid in ultrapure water with 0.1 %formic acid) and directed to the triple-quadrupole mass spectrometer (API 4000, Applied Biosystems, Foster City, CA, USA). The acquisition was run in multi-reaction mode (MRM) for 3-7 min. The following multi reaction monitoring (MRM) transitions were monitored; 349.3 ->261.0 for GABA-SymDaq complex and 355.1 -> 267.1 for GABA-d6-SymDaq complex, 393.3 -> 305.1 for glutamate-SymDaQ complex, 398.3 ->310.1 for glutamate-d5-SymDaQ complex. The absolute concentrations of GABA and glutamate were calculated using Analyst 1.4.2 software (Applied biosystems, Foster City, CA, USA), using analyte/IS ratio versus concentration, the calibrations curve was linear with 1/x weighing function. Lower limit of quantitation was 0.8 nM on column for GABA and 0.02 µM on column for glutamate and inter-day variation were 6.68 %CV for GABA and 3.77 %CV for glutamate.

### 4. Experiment 2: Effects of pharmacological manipulations of glutamate and GABA signaling in the LS on social play behavior in male and female juvenile rats

To determine the causal involvement of the observed changes in glutamate and GABA release in the LS during social play and in response to a V1aR antagonist, the effects of drugs targeting glutamate receptors (NMDA, kainate, AMPA) or the GABA-A receptor in the absence or presence of a V1aR antagonist were measured on social play behavior in male and female juvenile rats.

#### 4.1. Cannulation surgery

After five days of daily handling, male and female juvenile rats (30 days of age), were anesthetized with isoflurane (Butler Schein Animal Health, Dublin, OH) and mounted on a stereotaxic frame with the tooth bar set at − 4.5 mm. Guide cannulae (22 gauge; Plastics One, Roanoke, VA) were implanted 2 mm dorsal to the medial part of the LS according to Veenema et al. (2013) at coordinates 0.4 mm caudal to bregma, − 1.0 mm lateral to the midline and 3.6 mm ventral to the surface of the skull under an angle of 10° from the midsagittal plane to avoid damage to the sagittal sinus. Cannulae were fixed to the skull with three stainless steel screws and dental cement and closed with a dummy cannula (Plastics One, Roanoke, VA). After surgery, rats were housed individually and exposed to the social play test two and three days later.

#### 4.2. Microinjection procedure

Male and female juvenile rats (32-33 days of age) received a single 0.5 µl injection of Ringer’s solution (vehicle) or one of the following drugs or cocktails: the non-specific glutamate receptor agonist L-glutamic acid (600 nmol; Sigma Aldrich, St. Louis, MO), the V1aR antagonist (CH_2_)_5_Tyr(Me^2^)AVP (10ng), a cocktail containing the V1aR antagonist d(CH_2_)_5_Tyr(Me^2^)AVP (10ng) and L-glutamic acid (600nmol), a cocktail containing the V1aR antagonist d(CH_2_)_5_Tyr(Me^2^)AVP (10ng), NMDA receptor antagonist AP-5 (2 mM; Sigma Aldrich, St. Louis, MO), and AMPA receptor antagonist CNQX (0.4 mM; Sigma Aldrich, St. Louis, MO), or the selective GABA-A receptor agonist muscimol (100 ng; Sigma Aldrich, St. Louis, MO). Each rat received one vehicle injection and one drug injection in a counterbalanced order over two consecutive days. Drug doses were chosen based on previous microinjection studies in rats inducing changes in behavior (Veenema et al., 2012, 2013; Bredewold et al., 2015; Stanley et al., 1993; Numan et al., 2010).

#### 4.3. Histological verification of cannula placement

After the experiments, rats were killed with CO_2_, and charcoal was injected into their brains as a marker to check proper cannula placement histologically on Nissl-stained cryocut coronal brain sections. Rats with incorrect cannula placement were removed from the analysis.

### 5. Experiment 3. Analysis of V1aR mRNA-expressing cell types in the LS of male and female juvenile rats

To determine potential sex differences in V1aR mRNA-expressing cell types, fluorescent in situ hybridization was performed detecting expression of Avpr1a on GABAergic neurons (using Gad1 and Gad2 as markers), GABAergic interneurons [using somatostatin (Sst) or calbindin 1 (Calb1) as markers] and on astrocytes (using Sox9 as marker).

#### 5.1. Fluorescent in situ hybridization

Juvenile male and female rats (32-33 days of age, n=5- 7/sex) were euthanized with CO_2_, and brains were flash frozen on n-methylbutane, kept on dry ice, and stored at –80°C. Brains were blocked for the LS and 16 μm coronal LS sections were cut on the cryostat and mounted on SuperFrost Plus slides. Slides were stored at –80°C. RNAscope V2 kits (Advanced Cell Diagnostics, Newark, CA, USA) and probes to detect Avpr1a, Gad1, Gad2, Sst, Calb1, and Sox9 were used according to the manufacturer’s instructions. Tissue sections were washed in 1x phosphate-buffered saline (pH 7.6), dried at 60^0^C for 30 min, post-fixed in 4% paraformaldehyde in 1x phosphate-buffered saline for 1 h at 4°C, and dehydrated in 50% ethanol (1 x 5 min), 70% ethanol (1 x 5 min), and 100% ethanol (2 x 5 min). Following a 10- minute hydrogen peroxide incubation and target retrieval in a steamer, tissue sections were treated with protease III for 30 min at RT. Consecutive sets of sections were probed for Avpr1a/Gad1/Gad2, Avpr1a/Sst, Avpr1a/Calb1, and Avpr1a/Sox9. The probes were diluted 1:50 and applied to slides for 2 h at 40° C. Next, slides were rinsed in wash buffer and incubated in amplification buffer (AMP) 1 for 30 min at 40° C, then rinsed again in wash buffer and incubated in AMP 2 for 30 min at 40° C, then rinsed again in wash buffer and incubated in AMP 3 for 15 min at 40° C. Finally, slides were developed for 30 min in GFP (Gad1, Sst, and Avpr1a for run with Calb1), Cy3 (Gad2, Sox9, Calb1), and Cy5 (Avpr1a for runs with Gad1, Gad2, Sst and Sox9). Slides were washed in wash buffer and coverslipped with Vectashield hardset antifade mounting medium with a DAPI counterstain (Vector Laboratories, Burlingame, CA) and stored at 4°C.

#### 5.2 Image acquisition and colocalization analyses

Images were acquired with a 20x objective on a BZ-X700E/BZ-X710 fluorescence microscope and the associated BZH3AE software (Keyence Corporation of America, Elmwood Park, NJ, USA). Five bilateral sets of images were taken for both the dorsal and intermediate LS from bregma 0.84 – 0.36 mm (Paxinos & Watson, 2007; Fig 4A). Counts were conducted for the entirety of each image obtained (image field-of-view = 724.69 × 543.52 μm). Nuclei with ≥ 3 puncta positive for Avpr1a were counted and then determined to be positive (≥ 3 puncta) or negative (:< 2 puncta) for Gad1+Gad2, Sst, Calb1, or Sox9.

### 6. Statistics

For experiment 1, absolute levels of glutamate and GABA as well as glutamate/GABA concentration ratios (glutamate concentration in nM/GABA concentration in nM) were analyzed using one-way ANOVAs with sex as factor. Percentages of glutamate and GABA release were analyzed using three-way ANOVAs for repeated measures (sex x drug x dialysate). Interaction effects were followed by LSD post-hoc tests. When main effects for dialysate were found, one-way ANOVAs for repeated measures were run. For experiment 2, effects of drug treatment on social play behavior were analyzed using two-way ANOVAs (sex x drug). When interaction effects were found, Bonferroni post-hoc tests were run to test for differences between groups. When only a main effect of drug treatment was found, one-way ANOVAs were run separately for sex to test for differences between treatments. For experiment 3, the number of Avpr1a+ cells and the percentages of Avpr1a+ cells co-expressing Gad1/Gad2, Sst, Calb1, and Sox9 were analyzed using one-way ANOVAs with sex as factor. Data are presented as mean + SEM. Significance was set at p < 0.05.

## RESULTS

### Experiment 1. Effects of LS-V1aR blockade on extracellular glutamate and GABA release in the LS under baseline conditions and during exposure to social play in male and female juvenile rats

*Extracellular glutamate concentrations*. Males showed higher extracellular glutamate concentrations compared to females under baseline condition (p < 0.05; Fig 1A, B) and during social play (p< 0.05; Fig. 1A). V1aR antagonist treatment eliminated these sex differences under baseline condition (p = 0.42) and during social play (p = 0.21; Fig. 1B).

**Fig. 1.**
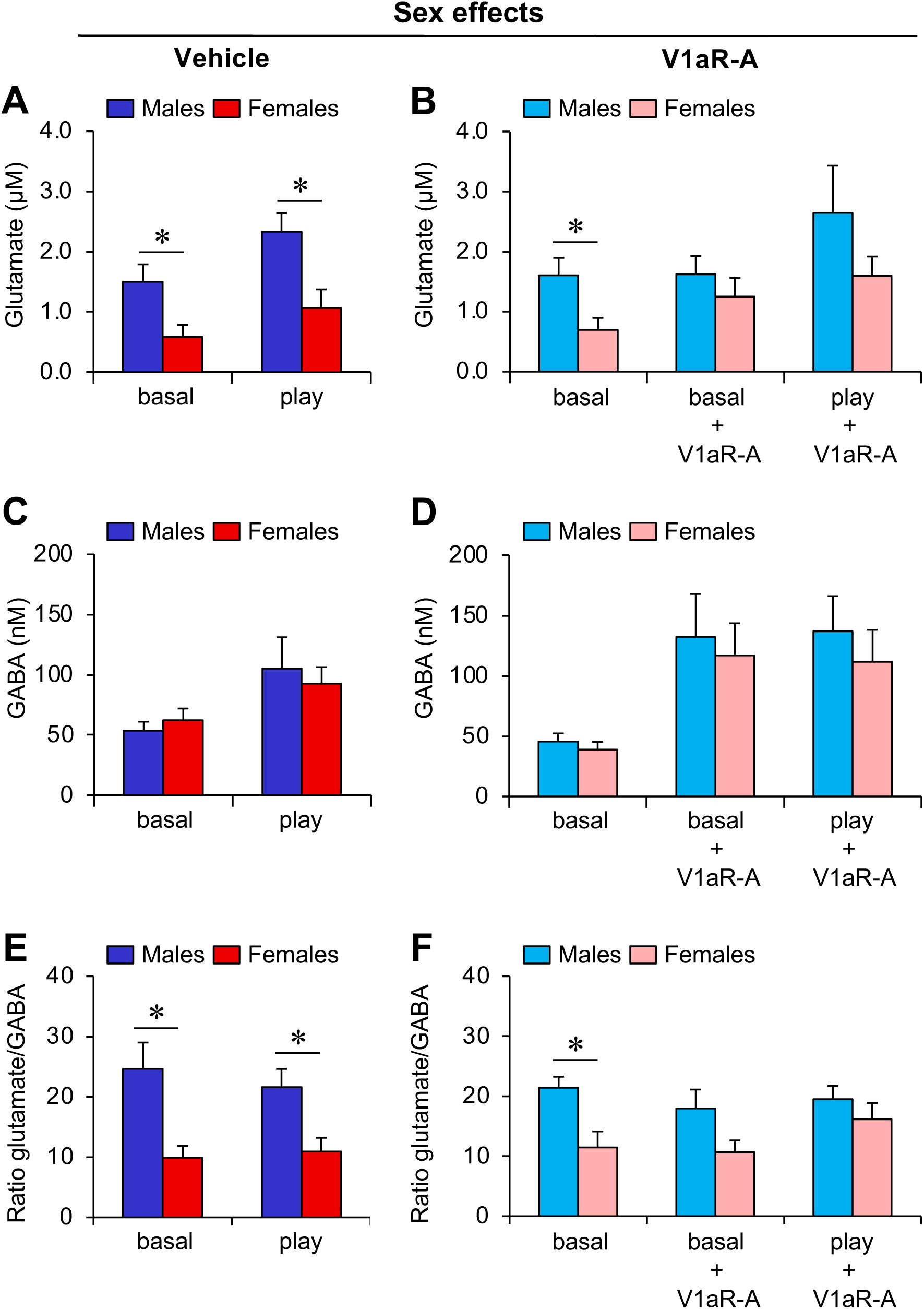
Sex differences in extracellular glutamate concentration and in glutamate/GABA ratio in the LS of juvenile rats are eliminated by LS-V1aR blockade. **(A-B)** Extracellular glutamate concentration is higher in males than in females under baseline conditions (basal; **A-B**) and during social play (play; **B**); This effect is abolished by V1aR antagonist (V1aR-A) administration into the LS (**B**). (**C-D**) Males and females show similar extracellular GABA concentrations independent of social play exposure (**C**) or V1aR-A treatment (**D**). (**E-F**) The glutamate/GABA ratio (glutamate concentration in nM/GABA concentration in nM) is higher in males than in females under baseline conditions and during social play (**E**); This effect is abolished by V1aR-A administration into the LS (**F**). Data represent mean + SEM. *p < 0.05, one-way ANOVA.

*Extracellular GABA concentrations.* No significant sex differences were found in extracellular GABA concentrations under baseline condition or during exposure to the social play test with or without V1aR antagonist treatment (p;= 0.49; Fig. 1C, 1D).

*Glutamate/GABA concentration ratio.* Males have a higher glutamate/GABA concentration ratio compared to females under baseline condition and during exposure to the social play test (p < 0.05; Fig. 1E). V1aR antagonist treatment abolishes the sex difference in glutamate/GABA concentration ratio (V1aR-A: p = 0.06; V1aR-A + play: p =0.36; Fig. 1F).

*Percentage glutamate release*. In males, V1aR antagonist treatment did not change the percentage release of glutamate (Treatment effect: F_(1,15)_ = 0.15; p = 0.71; Fig. 2A). However, exposure to the social play test was associated with an increase in the percentage glutamate release (Dialysate effect: F_(5,75)_ = 5.55; p < 0.001; Fig. 2A). Subsequent one-way ANOVAs for repeated measures revealed that exposure to the social play test was associated with a significant increase in the percentage glutamate release in vehicle-treated males (F_(5,35)_ = 7.94; p < 0.001; Dialysate 4 versus all other dialysates: p < 0.05) and V1aR antagonist-treated males (F_(5,35)_ = 6.24; p < 0.001; Dialysate versus dialysate 1-3, and 6: p < 0.05; Fig. 2A).

**Fig. 2.**
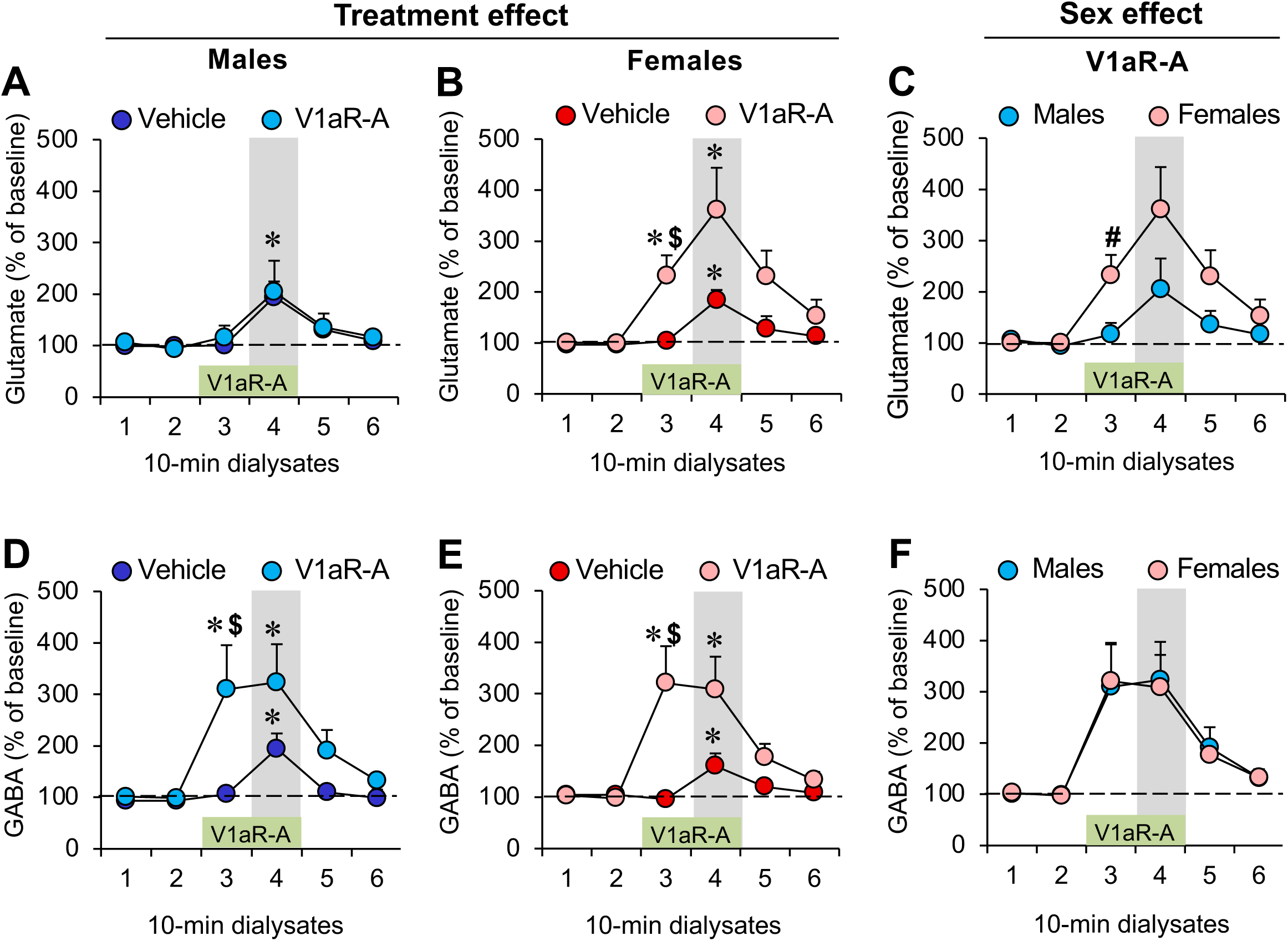
Sex-specific increase in extracellular glutamate release, but not GABA release, in the lateral septum (LS) of juvenile rats in response to LS-V1aR blockade. **(A-B)** Extracellular glutamate release is increased during social play (dialysate 4, grey bar) in both males (**A**) and females (**B**); V1aR antagonist (V1aR-A) administration into the LS (during dialysate 3 and 4, green bar) increased glutamate release prior to the social play test (dialysate 3) in females (**B**) but not in males (**C**) Comparing males and females shows that V1aR-A administration into the LS induces higher glutamate release prior to the social play test in females versus males. (**D-F**) Males and females show similar increases in extracellular GABA release during social play and in response to V1aR-A administration into the LS. Data represent mean + SEM. *p < 0.05 versus baseline (dialysate 1 and 2), $ p < 0.05 versus vehicle (**B, D, E**), # p < 0.05 versus males (**C**); two-way ANOVA for repeated measures followed by one-way ANOVA for repeated measures and/or LSD post hoc test.

In females, V1aR antagonist treatment increased the percentage release of glutamate (Dialysate x Treatment effect: F_(5,80)_ = 2.48; p < 0.05) prior to exposure to the social play test (dialysate 3) compared to baseline (dialysates 1 and 2; p < 0.01) and compared to vehicle treatment (dialysate 3: p < 0.05; Fig. 2B). Furthermore, exposure to the social play test (dialysate 4) was associated with a significant increase in the percentage glutamate release compared to baseline in both vehicle-treated females (dialysates 1-3: p < 0.01) and V1aR antagonist-treated females (dialysates 1 and 2: p < 0.05; Fig. 2B).

Lastly, V1aR treatment resulted in a significantly higher percentage glutamate release in females compared to males (Sex effect: F_(1,17)_ = 4.64; p < 0.05) prior to exposure to the social play test (dialysate 3; p < 0.05; Fig. 2C).

*Percentage GABA release.* In males, V1aR antagonist treatment increased the percentage release of GABA (Dialysate X Treatment effect: F_(5,75)_ = 8.23; p < 0.05) prior to exposure to the social play test (dialysate 3) compared to dialysates 1 and 2 (p < 0.01) and compared to vehicle treatment (dialysate 3: p < 0.05; Fig. 2D). Furthermore, exposure to the social play test was associated with an increase in the percentage GABA release compared to baseline in both vehicle-treated males (dialysates 1-3: p < 0.05) and V1aR antagonist-treated males (dialysates 1 and 2: p < 0.05; Fig. 2D).

In females, V1aR antagonist treatment increased the percentage release of GABA (Dialysate x Treatment effect: F_(5,80)_ = 4.65; p < 0.005) prior to exposure to the social play test (dialysate 3) compared to dialysates 1 and 2 (p < 0.005) and compared to vehicle treatment (dialysate 3: p < 0.05; Fig. 2E). Furthermore, exposure to the social play test (dialysate 4) was associated with a significant increase in the percentage GABA release compared to baseline in both vehicle-treated females (dialysate 3: p < 0.05) and V1aR antagonist-treated females (dialysates 1 and 2: p < 0.05; Fig. 2E).

Lastly, V1aR treatment resulted in similar increases in percentage glutamate release in males compared to females (Sex effect: F_(1,17)_ = 0.007; p = 0.93) prior and during exposure to the social play test (Fig 2F).

### Experiment 2. Effects of pharmacological manipulations of glutamate and GABA neurotransmission in the LS on social play behavior in male and female juvenile rats

We first determined whether and how the V1aR antagonist-induced increase in glutamate release observed in females only is involved in the regulation of social play behavior. To this end, male and female juvenile rats received microinfusions into the LS with either a vehicle solution or L-glutamic acid into the LS 20 min prior to exposure to the social play test. L-glutamic acid reduced the duration of social play in both sexes (Treatment effect: F_(1,32)_ = 22.4; p < 0.001; males, p < 0.01; females, p < 0.001; Fig. 3A). Furthermore, there was a sex effect on social play duration (Sex effect: F_(1,32)_ = 4.56; p < 0.05) which is driven by L-glutamic acid inducing a shorter duration of social play in females compared to males (p < 0.05; Fig. 3A).

**Fig. 3.**
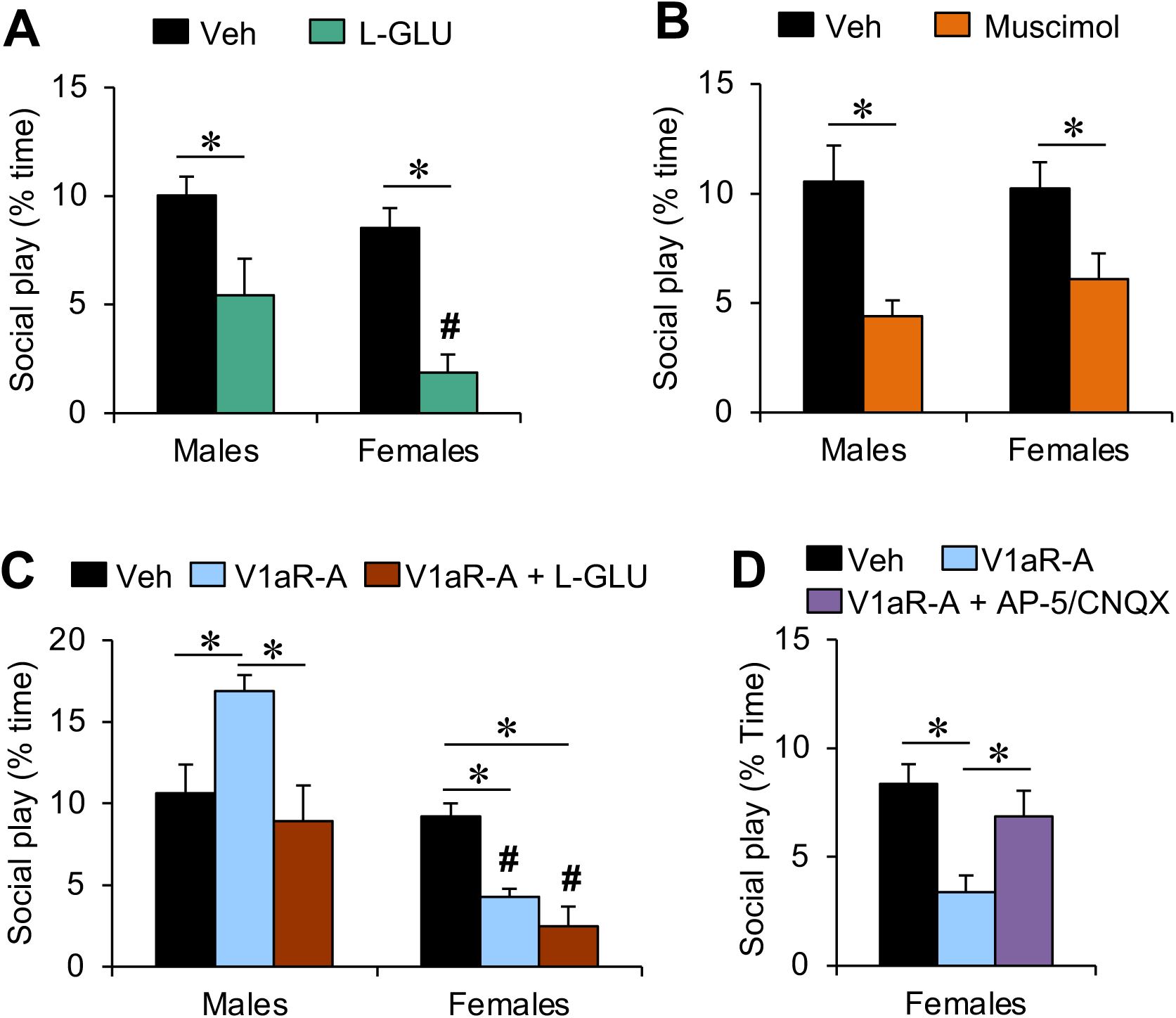
Effects of pharmacological manipulations of glutamate and GABA neurotransmission in the LS on social play behavior in male and female juvenile rats. **(A)** The non-specific glutamate receptor agonist L-glutamic acid (L-GLU; 600 nM/0.5 ml) decreased social play duration in both sexes, with a stronger effect in females compared with males. (**B**) The GABA-A receptor agonist muscimol (100 ng/0.5 ml) decreased social play duration in both sexes. (**C**) Co-administration of L-GLU (600 nmol) with the V1aR antagonist (V1aR-A; 10ng/0.5 ml) prevented the V1aR-A-induced increase in social play duration in males while it did not change the V1aR-A-induced decrease in social play duration in females. (**D**) Co-administration of the NMDA receptor antagonist AP-5 and the AMPA/kainate receptor antagonist CNQX (2 mM+0.4 mM/0.5 ml) prevented the V1aR-A-induced decrease in social play duration in females. The duration of social play behavior is expressed as percentage of total time. Data represent mean + SEM. *p < 0.05 vs. vehicle, # p < 0.05 vs. males, two-way ANOVA followed either by one-way ANOVA (**A, B**) or Bonferroni post hoc test (**C, D**). Veh, vehicle.

Next, we determined whether and how the V1aR antagonist-induced increase in GABA release observed in both sexes is involved in the regulation of social play behavior. Accordingly, male and female juvenile rats received microinfusions into the LS with either a vehicle solution or the GABA-A receptor agonist muscimol into the LS 20 min prior to exposure to the social play test. Administration of muscimol reduced social play duration in both sexes (Treatment effect: F_(1,26)_ = 17.3; p < 0.001; males: p < 0.005; females: p < 0.05; Fig. 3B).

We then determined whether an increase in glutamate neurotransmission would prevent the V1aR antagonist-induced increase in social play levels in males. L-glutamic acid was administered into the LS in conjunction with the V1aR antagonist 20 min prior to exposure to the social play test. A sex x treatment effect was found for the duration of social play (Sex x Treatment effect: F_(2,46)_ = 8.43; p < 0.005). Bonferroni posthoc testing revealed that, in males, the V1aR antagonist-induced increase in social play duration (V1aR-A vs vehicle: p < 0.01) is abolished by co-administration of L-glutamic acid (V1aR-A + L-GLU vs V1aR-A: p < 0.005; V1aR-A + L-GLU vs vehicle: p = 1.00; Fig. 3C). In females, V1aR antagonist (p < 0.05) and the combined treatment of the V1aR antagonist and L-glutamic acid (p < 0.005) reduced social play duration compared to vehicle (Fig. 3C). This resulted in a sex difference in social play duration in both treatment groups with shorter social play duration in females compared to males (V1aR-A: p < 0.005; V1aR-A + L-GLU: p < 0.001; Fig. 3C).

Lastly, we determined whether administration of the ionotropic glutamate receptor antagonists AP-5 and CNQX into the LS at a dose (2mM+0.4 mM/0.5 μl) that did not alter social play duration (Bredewold et al., 2015), would prevent the previously observed decrease in social play duration induced by V1aR antagonist administration in the LS of female juvenile rats (Bredewold et al., 2015; Veenema et al., 2013). A main effect of treatment was found for social play duration (Treatment effect: F_(1,24)_ = 5.28; p < 0.05). Bonferroni posthoc testing revealed that V1aR antagonist administration into the LS reduced social play duration (V1aR-A vs vehicle: p < 0.05) and this effect was abolished by simultaneous administration of AP-5 and CNQX (AP- 5+CNQX vs vehicle: p = 1.00; AP-5/CNQX vs V1aR-A: p < 0.05; Fig 3D).

### Experiment 3. Analysis of Avpr1a-expressing cell types in the LS of male and female juvenile rats

We determined potential sex differences in the number of Avpr1a+ cells and in the percentage of Avpr1a-expressing cell types in the LS. No sex differences were found in the number of Avpr1a-expressing cells in the dorsal LS or intermediate LS (Table 1). Likewise, no sex differences were found in the percentages of Avpr1a-expressing cells in the dorsal LS (Fig. 4) or intermediate LS (Table 2) co-expressing Gad1/Gad2, Sst, Calb1, or Sox9.

**Fig. 4.**
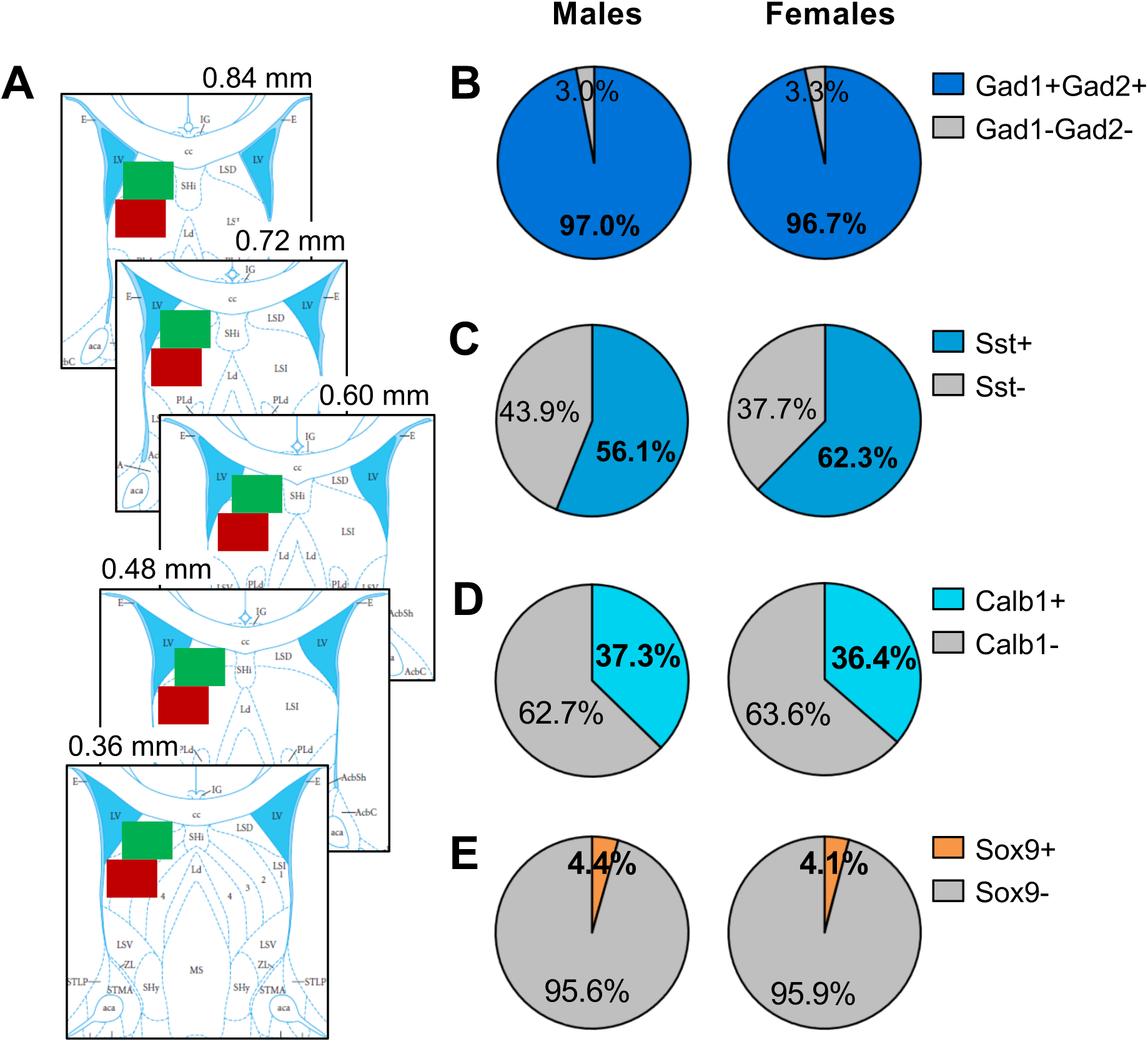
No sex differences in the percentage of Avpr1a+ cells expressing markers for GABAergic neurons, somatostatin-expressing GABAergic neurons, calbindin-expressing GABAergic neurons or astrocytes in the dorsal LS of juvenile rats. **(A)** Example image acquisition locations (showing unilateral for simplicity) for dorsal LS (green) and intermediate LS (red) on rat brain atlas templates (bregma 0.84 – 0.36 mm; Paxinos & Watson, 2007) (**B-E**) There were no sex differences in the percentage of Avpr1a+ cells in the dorsal LS co-expressing Gad1 and Gad2 (**B**), co-expressing Sst (**C**), co-expressing Calb1 (**D**), or co-expressing Sox9 (**E**). Data represent mean; one-way ANOVA.

**Table 1.**
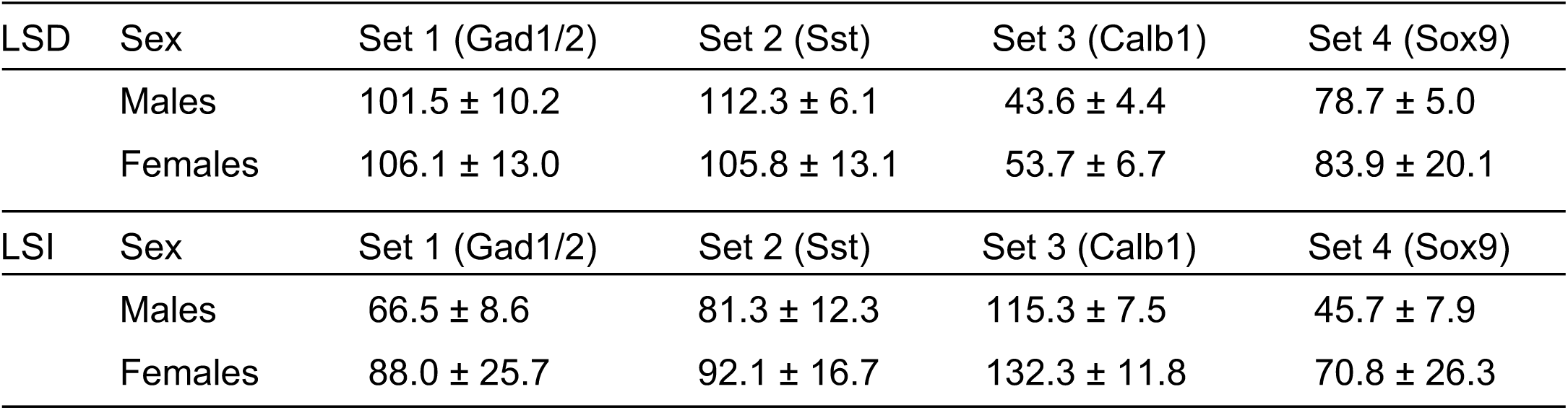
No sex differences in the number of Avpr1a+ cells in the LS of juvenile rats. Using fluorescent in situ hybridization, Avpr1a+ cells were counted in the dorsal LS (LSD) and intermediate LS (LSI) of male and female juvenile rats (32-33 days of age; n=4-7/sex). Cells were considered positive for Avpr1a when expressing ≥ 3 puncta per cell. Males and females had similar numbers of Avpr1a+ cells across Sets 1 - 4. Sets 1 - 4 refers to the LS section sets that were probed for mRNA transcripts for Avpr1a along with either Gad1/2 (Set 1), somatostatin (Sst, Set 2), calbindin (Calb1; Set 3) or Sox9 (Set 4). Data represent mean ± SEM; One-way ANOVA.

| LSD | Sex | Set 1 (Gad1/2) | Set 2 (Sst) | Set 3 (Calb1) | Set 4 (Sox9) |
| --- | --- | --- | --- | --- | --- |
| | Males | 101.5 ± 10.2 | 112.3 ± 6.1 | 43.6 ± 4.4 | 78.7 ± 5.0 |
| | Females | 106.1 ± 13.0 | 105.8 ± 13.1 | 53.7 ± 6.7 | 83.9 ± 20.1 |
| LSI | Sex | Set 1 (Gad1/2) | Set 2 (Sst) | Set 3 (Calb1) | Set 4 (Sox9) |
| | Males | 66.5 ± 8.6 | 81.3 ± 12.3 | 115.3 ± 7.5 | 45.7 ± 7.9 |
| | Females | 88.0 ± 25.7 | 92.1 ± 16.7 | 132.3 ± 11.8 | 70.8 ± 26.3 |

**Table 2.** No sex differences in the percentage of Avpr1a+ cells co-expressing markers for either GABAergic neurons, somatostatin-expressing GABAergic neurons, calbindin-expressing GABAergic neurons or astrocytes in the intermediate LS of juvenile rats. Percentage of Avpr1a+ cells in the intermediate LS co-expressing Gad1 and Gad2 (markers for GABAergic neurons), co-expressing somatostatin (Sst), co-expressing calbindin (Calb1), or co-expressing Sox9 (marker for astrocytes) did not differ between the sexes. Cells were considered positive for a marker when expressing ζ 3 puncta per cell. Data represent mean ± SEM; One-way ANOVA.

| Sex | Gad1/2 | Sst | Calb1 | Sox9 |
| --- | --- | --- | --- | --- |
| Males | 95.5 ± 1.3 | 41.9 ± 4.2 | 28.2 ± 2.6 | 6.9 ± 0.9 |
| Females | 96.3 ± 0.5 | 46.5 ± 4.2 | 27.4 ± 3.7 | 6.0 ± 1.0 |

## DISCUSSION

We previously showed that the LS-AVP system regulates social play behavior in opposite directions in male and female juvenile rats (Veenema et al., 2013; Bredewold et al., 2014). We now demonstrate a potential role for glutamate, but not for GABA, in this sex-specific regulation of social play by the LS-AVP system. We first showed that V1aR antagonist administration into the LS increased the extracellular release of glutamate in the LS of females, but not of males. No sex difference were found in extracellular GABA release in the LS. Accordingly, the baseline sex differences in extracellular glutamate concentration and in glutamate/GABA ratio (both higher in males) were eliminated by LS-V1aR blockade. Second, we found that administration of L-glutamic acid into the LS decreased social play behavior and prevented the V1aR antagonist-induced increase in social play behavior in males. Third, we found that administration of L-glutamic acid into the LS mimicked the V1aR antagonist-induced decrease in social play behavior in females. Fourth, we showed in females that antagonizing glutamate receptors prevented the V1aR antagonist-induced decrease in social play behavior. Together, these findings suggest that the V1aR antagonist-induced decrease in social play behavior in females is mediated by enhanced LS-glutamate signaling. In contrast, the mechanism by which LS-V1aR blockade increased social play behavior in males seems independent of LS-glutamate signaling. Last, we found no sex differences in different types of GABAergic neurons expressing the V1aR nor in V1aR-expressing astrocytes in the LS. This suggests that the cellular mechanism by which LS-V1aR activation sex-specifically modulates LS- glutamate release and social play behavior may not involve LS-GABAergic neurons or LS- astrocytes.

We confirmed previous findings that males have higher extracellular glutamate, but not GABA, concentrations in the LS compared to females, resulting in a higher glutamate/GABA ratio in males under baseline conditions as well as during exposure to the social play test (Bredewold et al., 2015). Despite this sex difference, males and females show similar levels of social play behavior in our testing conditions (current study; Veenema et al., 2013; Bredewold et al., 2014, 2015, 2018; Reppucci et al., 2018, 2020). We now show that LS-V1aR blockade abolished the sex difference in extracellular glutamate concentrations as well as in glutamate/GABA ratio under baseline conditions and during exposure to the social play test. Furthermore, we confirmed previous findings that LS-V1aR blockade induced a sex difference in social play (Veenema et al., 2013; Bredewold et al., 2015). Taken together, these findings strongly suggest that by making the sexes more alike (i.e., similar glutamate concentrations and glutamate/GABA ratio), a sex difference in social play behavior is induced. Although perhaps a surprising finding, it is in line with the compensation theory of sex differences proposed by De Vries and colleagues (De Vries & Boyle, 1998; De Vries, 2004). This theory postulates that some sex differences in the brain may serve to prevent sex differences in behavior allowing males and females to display similar behaviors despite major sex differences in physiological and hormonal conditions. Thus, it would be interesting to explore whether the observed sex difference in glutamate/GABA ratio occurs across the lifespan, in additional brain regions, and in other species. In humans, a higher excitatory/inhibitory ratio is thought to contribute to the pathophysiology of various neurological and psychiatric disorders, including ASD (Nelson & Valakh, 2015; Rubenstein & Merzenich, 2003; Robertson et al., 2016). This contrasts with another study showing that a higher excitation/inhibition ratio during development is linked to neuroplasticity and the acquisition of new cognitive skills (Cohen Kadosh et al., 2015). Interestingly, the former studies are primarily performed in male subjects, while the latter used females only. Thus, our findings in rats may provide incentive to study possible sex differences in excitatory/inhibitory ratio in humans.

The observed female-specific increase in extracellular LS-glutamate release upon LS-V1aR blockade likely resulted in altered LS neuronal excitability, which in turn, may have contributed to the decrease in social play seen after LS-V1aR blockade in females. Indeed, we found that mimicking the increase in glutamate by administration of L-glutamic acid into the LS decreased social play in females. Furthermore, we showed that antagonizing ionotropic glutamate receptors (at a dose that did not change social play behavior in females; Bredewold et al., 2015) prevented the LS-V1aR blockade-induced decrease in social play behavior in females. This strongly suggests that glutamate is a downstream substrate by which LS-V1aR blockade reduced social play behavior in females, an effect that can be mimicked in males. In contrast to females, LS-V1aR blockade did not induce a change in extracellular glutamate release in males. However, L-glutamic acid administration into the LS induced a decrease in social play behavior in males, an effect seen in females too, and prevented the increase in social play behavior induced by LS-V1aR blockade in males. This may indicate that the mechanism by which enhanced glutamate neurotransmission in the LS decreases social play behavior is the same in males and females.

*In vitro* studies have shown that AVP via V1aR activation modulates glutamatergic synaptic currents in the LS (Joels & Urban, 1982, 1984, 1985) and directly excites LS GABAergic interneurons (Allaman-Exertier et al., 2007). These findings support the possibility that activation of V1aRs postsynaptically expressed on LS-GABAergic interneurons, can directly modulate glutamate release in the LS. Additionally, potential sex differences herein could explain the observed sex differences in extracellular LS-glutamate release under baseline conditions and upon LS-V1aR blockade. We showed co-expression of Avpr1a with Gad1 and Gad2, Sst, and Calb1. However, we found no sex differences in the percentage of Avpr1a+ cells co-expressing any of these markers in either the dorsal of intermediate LS. Notably, Sst-expressing cells (>50% in dorsal LS; >40% in intermediate LS) and Calb1-expressing cells (almost 40% in dorsal LS; almost 30% in intermediate LS) represent most Avpr1a-expressing cells in the LS, suggesting important roles for both subtypes of GABAergic interneurons in V1aR-mediated functions in the LS. Enhanced activity of Sst-expressing neurons in the dorsal LS is associated with increased anxiety-like behaviors (Besnard et al., 2019; An et al., 2022) while suppressed activity of Sst-expressing neurons in the dorsal LS is associated with improved social discrimination behavior (Borie et al., 2021) in mice. Calb1 is almost exclusively expressed in GABAergic neurons in the LS (Zhao et al., 2013), but the function of Calb1-expressing cells in the LS seems unknown. For future research, it would be interesting to specifically target these subtypes of GABAergic interneurons to determine their role in social play behavior in male and female rats.

Glutamate release and glutamate re-uptake in the brain are both regulated by astrocytes (Mahmoud et al., 2019). Thus, extracellular glutamate concentrations could be regulated through activation of V1aR expressed on astrocytes in the LS. Furthermore, a sex difference in V1aR-expressing astrocytes could contribute to a sex difference in extracellular LS-glutamate concentrations under baseline conditions and/or upon LS-V1aR blockade. We indeed demonstrated that a small percentage of Avpr1a+ cells co-express Sox9, which is a reliable marker of astrocytes (Sun et al., 2017), in the dorsal LS (around 4%) and intermediate LS (around 6%). However, there was no sex difference in the percentage of Avpr1a+ cells co-expressing Sox9. These findings suggest the potential of a source other than local astrocytes and local GABAergic neurons contributing to the observed sex differences in LS-glutamate release.

Extracellular glutamate released in the LS likely derives from glutamatergic neurons projecting to the LS, including those located in the hippocampus (Zaczek et al., 1979; Gallagher et al., 1995) and in the hypothalamus (Chee et al., 2015). Thus, a sex difference in extracellular LS- glutamate concentrations under baseline conditions and/or upon LS-V1aR blockade could be caused by activation of V1aR presynaptically expressed on glutamatergic afferents of the LS. Although this has not been described for the V1aR in the LS, *in vitro* studies have suggested that AVP acts on presynaptically expressed V1aR to modulate glutamate release in other brain regions, namely the parabrachial nucleus and the nucleus of the solitary tract (Chen & Pittman, 1999; Bailey et al., 2006). Interestingly, bath application of a V1aR antagonist increased glutamate release from solitary tract neurons (Bailey et al., 2006), suggesting that presynaptic V1aR activation leads to an inhibition of glutamate release. Future studies are required to determine whether V1aR is presynaptically expressed on glutamatergic terminals in the LS and whether there are sex differences in the presence and/or function of V1aR at these presynaptic glutamatergic terminals.

We found that LS-V1aR blockade increased extracellular LS-GABA release in both sexes. Because LS-V1aR blockade changed social play behavior in opposite directions in males versus females, it could be that enhanced LS-GABA signaling is the downstream substrate. However, we found that muscimol-induced activation of GABA-A receptors in the LS decreased social play behavior in both sexes. Interestingly, we previously showed that bicuculline-induced blockade of GABA-A receptors in the LS also decreased social play behaviors in both sexes (Bredewold et al., 2015). This suggests that any change in GABA-A receptor activation results in a decrease in social play behavior. GABA-A receptor activation presumably will result in decreased LS neuronal output while GABA-A receptor blockade will allow for increased LS output (Sheehan et al., 2004). It may therefore be surprising that changing GABAergic tone in either direction would have a similar effect on social play behavior. Yet, similar effects of bicuculline and muscimol on neuronal activation have been reported before (Yoon et al., 2010). Independent of the underlying mechanisms, our findings provide evidence indicating that although GABA neurotransmission in the LS modulates social play behavior in both sexes, it is not involved in the sex-specific regulation of social play behavior by the LS-AVP system.

In conclusion, our data demonstrate a role for glutamate as a downstream substrate by which the LS-AVP system regulates social play behavior in sex-specific ways. Further research is required to determine the cellular mechanism by which LS-V1aR activation alters LS-glutamate release and how this regulates social play behavior in sex-specific ways. We hypothesize that this may be through V1aRs presynaptically expressed on glutamatergic neurons projecting to the LS. Given the rewarding nature of social play behavior and its critical role in the development of social competence, understanding how social play behavior is regulated in sex-specific ways by the LS-AVP system will be crucial to our understanding of social play deficits in sex-biased neurodevelopmental disorders, such as ASD.

## ACKNOWLEDGEMENTS

We would like to thank Jennifer Schiavo, Michelle Verreij, and Nara Nascimento for technical assistance, members of the Veenema lab for input and critically reading the manuscript, Dr. Maurice Manning (Toledo, OH) for kindly providing the V1aR antagonists, and animal caretakers for excellent animal care. This research was supported by NARSAD Grant 17382, NSF IOS1253386, and NIH R01MH102456 to AHV.

